# Engineering Insulin Cold Chain Resilience to Improve Global Access

**DOI:** 10.1101/2021.04.13.439582

**Authors:** Caitlin L. Maikawa, Joseph L. Mann, Aadithya Kannan, Catherine M. Meis, Abigail K. Grosskopf, Ben S. Ou, Anton A. A. Smith, Gerald G. Fuller, David M. Maahs, Eric A. Appel

**Author notes:** These authors contributed equally (Caitlin L. Maikawa, Joseph L. Mann). Department of Health Technology, Technical University of Denmark, 2800 Kgs. Lyngby, Denmark. Person to whom correspondence should be addressed: Dr. Eric A. Appel.

## Abstract

There are 150 million people with diabetes worldwide who require insulin replacement therapy and the prevalence of diabetes is rising fastest in middle and low-income countries. Current formulations require costly refrigerated transport and storage to prevent loss of insulin integrity. This study shows the development of simple “drop-in” amphiphilic copolymer excipients to maintain formulation integrity, bioactivity, pharmacokinetics and pharmacodynamics for over 6 months when subjected to severe stressed aging conditions that cause current commercial formulation to fail in under 2 weeks. Further, when these copolymers are added to Humulin R (Eli Lilly) in original commercial packaging they prevent insulin aggregation for up to 4 days at 50 °C compared to less than 1 day for Humulin R alone. These copolymers demonstrate promise as simple formulation additives to increase the cold chain resilience of commercial insulin formulations, thereby expanding global access to these critical drugs for treatment of diabetes.

## INTRODUCTION

There are over 150 million people with diabetes requiring insulin replacement therapy worldwide.^1^ The worldwide population of people with insulin-deficient diabetes is increasing at a rate of 3-5% annually, with the highest increases in prevalence occurring in warm regions such as Africa, the Western Pacific, and the Middle East. ^1–2^ Unfortunately, insulin is prone to irreversible aggregation when exposed to high temperatures and/or agitation and requires careful storage and refrigerated transport (the cold chain) to retain activity over its shelf life. Maintaining insulin integrity presents a challenge for the pharmaceutical industry, health care providers, and people with diabetes worldwide.^3–4^ Indeed, annual global costs for the refrigerated transport of all biopharmaceuticals globally exceed $15 billion (USD), and losses due to interruptions in the cold chain reach $35 billion (USD) annually.^4^

A primary driver of the loss of formulation integrity is the propensity of proteins to aggregate at hydrophobic interfaces when exposed to elevated temperatures. ^5^ In the case of insulin formulations, agitation and elevated temperatures – conditions common to worldwide transport and cold chain interruptions – increase interactions between partially unfolded insulin monomers adsorbed to interfaces, leading to nucleation of insulin amyloid fibrils.^6–9^

Recent studies in insulin stabilization have relied on covalent or non-covalent attachment of hydrophilic polymers directly to insulin or through encapsulation of insulin or insulin crystals in hydrogels.^10–15^ While these strategies have successfully shielded insulin from interfacial adsorption or insulin-insulin interactions and increased insulin stability, they can also lead to increased absorption times and longer circulation times in vivo. A more translatable stabilization method would utilize an inactive excipient that could be incorporated into existing formulations without altering the drug pharmacokinetics.

Amphiphilic copolymers present an alternative to polymer-protein conjugation, exploiting their propensity to gather at the air-water interface to hinder insulin-interface interactions.^6, 16–18^ Poloxamers have been successfully used to improve insulin stability and have been employed in commercial formulations (Insuman U400, Sanofi-Aventis). Yet, poloxamers have not been widely adopted and have not succeeded in reducing reliance on the cold chain for insulin transport.

Previously we have shown that biocompatible and non-toxic acrylamide carrier/dopant copolymers (AC/DC), a class of amphiphilic copolymers comprising water-soluble “carrier” and hydrophobic “dopant” monomers, can enable the development of ultra-fast insulin formulations.^19^ We hypothesized that these excipients could be applied more broadly to improve the stability of current commercial insulin formulations in the context of reducing cold chain reliance and improving formulation resilience. In this work, we aim to understand the stabilization mechanism of AC/DC copolymer excipients and to test the limits of their stabilizing capacity long-term and under extreme environmental conditions. We report the ability of select copolymers to act as simple “drop-in” excipients to stabilize commercial insulin formulations (Humulin R, Eli Lilly) without altering their bioactivity, pharmacokinetics or pharmacodynamics, constituting an important step toward improving global access to these critical drugs.

## EXPERIMENTAL METHODS

### Materials

Humulin R (Eli Lilly) was purchased and used as received. Solvents N,N-dimethylformamide (DMF; HPLC Grade, Alfa Aeser, >99.7%), hexanes (Fisher, Certified ACS, >99.9%), ether (Sigma, Certified ACS, Anhydrous,>99%) and CDCl_3_ (Acros, >99.8%) were used as received. Monomers N-(3-methoxypropyl)acrylamide (MPAM; Sigma, 95%), 4-acryloylmorpholine (MORPH; Sigma, >97%) were filtered with basic alumina prior to use. Monomers N-phenylacrylamide (PHE; Sigma, 99%) and N-isopropylacrylamide (NIPAM; Sigma, >99%) were used as received. RAFT chain transfer agents 2-cyano-2-propyl dodecyl trithiocarbonate (2-CPDT; Strem Chemicals, >97%) and 4-((((2-carboxyethyl)thio)carbonothioyl)thio)-4-cyanopentanoic acid (BM1433; Boron Molecular, >95%) were used as received. Initiator 2,2’-azobis(2-methyl-propionitrile) (AIBN; Sigma, >98%) was recrystallized from methanol (MeOH; Fisher, HPLC Grade, >99.9%) and dried under vacuum before use. Z-group removing agents lauroyl peroxide (LPO; Sigma, 97%) and hydrogen peroxide (H2O2; Sigma, 30%) were used as received. Streptozotocin (99.58%) was purchased from MedChem Express. All other reagents were purchased from Sigma-Aldrich unless otherwise specified.

### Surface Tension

Time resolved surface tension of the air-solution interface was measured with a Platinum/Iridium Wilhelmy plate connected to an electrobalance (KSV Nima, Finland). The Wilhelmy plate was partially immersed in the aqueous solution in a Petri dish, and the surface tension of the interface was recorded for 50 minutes from the formation of a fresh interface. Equilibrium surface tension values (t = 50 min) were reported as these values more closely describe the environment in a stored vial before agitation. Two replicates were taken and averaged.

### Interfacial Rheology

Interfacial shear rheology was measured using the Discovery HR-3 rheometer (TA Instruments) with an interfacial geometry comprising of a Du Noüy ring made of Platinum/Iridium wire (CSC Scientific, Fairfax, VA, catalog No. 70542000). Before each experiment, the Du Noüy ring was rinsed with ethanol and water and flame treated to remove organic contaminants. The solution chamber consisted of a double-wall Couette flow cell with an internal Teflon cylinder and an external glass beaker. A time-sweep was performed with a strain of 1% (within the linear regime) and a frequency of 0.05 Hz (low enough for instrument inertia to not be significant). Interfacial complex shear viscosity was measured for 30 minutes. The experiment was repeated in triplicate.

### Polymer Synthesis

Polymers were synthesized via reversible addition fragmentation transfer as described in previous literature. Detailed methods can be found in the supplementary information. Resulting composition and molecular weights were determined via ^1^H NMR spectroscopy and SEC with poly(ethylene glycol) standards (**Table S1**).

### In Vitro Insulin Stability Assay (Accelerated Aging)

50 μL of AC/DC excipient (MoNi_23%_, MpPhe_8%_, MoPhe_6%_) in milli-Q water (2.1, 21, or 105 mg/mL) or 50 μL of milli-Q water was added to 1 mL of Humulin R (Eli Lilly 100U) in a glass autosampler vial (J.G. Finneran, 2.0mL Clear R.A.M.™ Large Opening Vial, 12 x 32mm, 9mm Thread) and capped, yielding 95U Humulin either as a control or formulated with 0.01, 0.1, or 0.5 wt.% AC/DC excipient. These vials were incubated at 37 °C and agitated at 150 RPM for 2, 4, and 6 months (in addition, Humulin only control was agitated at 2 weeks and 1 month). The preparation of formulations was staggered so that all samples reached their endpoint age at the same time. Vials were refrigerated until testing upon reaching selected aging timepoint. Following initial transmittance experiments, all further experiments were done with formulations with 0.01 wt.% AC/DC excipient to minimize copolymer concentration. In addition, 500 μL of 2.1 mg/mL MoNi_23%_ or milli-Q water were added to 10 mL of unadulterated Humulin (Eli Lilly 100U) in its commercial vial to generate 95U Humulin control and formulation with 0.01 wt.% AC/DC excipient. These vials were placed in the original individual packaging boxes with the instruction papers. These packages were incubated at either 37 °C or 50 °C until significant opacity change. 300-400 μL aliquots were removed every 24 hours for the first 7 days and refrigerated. Following that, intermittent aliquots were taken to conserve volume. Every 24 hours, the bottoms of the vials were photographed to track the change in opacity. Methods for aggregation assays for recombinant human insulin were adapted from Webber *et al*.^11^ Formulation samples were plated at 150 μL per well in a clear 96-well plate and an absorbance reading was taken at 540 nm (BioTek Synergy H1 microplate reader). The aggregation of insulin leads to light scattering, which results in an increase in the measured absorbance. The time-to-aggregation (*t*_A_) was defined as the timepoint when a 10% increase in transmittance from time zero was observed.

### Circular dichroism

Circular dichroism was used to validate that aging with AC/DC excipients does not result in changes to the secondary structure of insulin. Aged Humulin (0.5, 1, 2, 4, and 6 months) or Humulin aged with 0.01 wt.% AC/DC excipients (2, 4, and 6 months) were evaluated against an unaged Humulin control or unaged Humulin with 0.01 wt.% AC/DC excipient. Formulation samples were diluted to 0.2 mg/mL in PBS (pH=7.4). Samples were left to equilibrate for 15 minutes at room temperature before measurement. Near-UV circular dichroism spectroscopy was performed at 20 °C with a J-815 CD Spectropolarimeter (Jasco Corporation) over a wavelength range of 200–260 nm using a 0.1 cm pathlength cell.

### In vitro insulin cellular activity assay

In vitro insulin activity was tested using the AKT phosphorylation pathway using AlphaLISA SureFire Ultra (Perkin-Elmer) kits for detection of phosphorylated AKT 1/2/3 (pS473) compared to total Akt1. Humulin, Aged Humulin (t = 6 months), Humulin + MoNi_23%_, and Aged Humulin + MoNi_23%_ (t = 6 months) formulations were tested. Methods have been previously described elsewhere.^13–14^ Detailed methods can be found in the supplemental information. Results were plotted as a ratio of [pAKT]/[AKT] for each sample (n=3 cellular replicates) and an EC_50_ regression (log(agonist) vs. response (three parameters)) was plotted using GraphPad Prism 8.

### In vitro cytotoxicity of AC/DC excipients

NIH 3T3 culture: NIH 3T3s were cultured according to ATCC’s recommendations. Media consisted of Dulbecco’s Modified Eagle Medium, 10% FBS and 1% Penicillin-Streptomycin. Cells were split at an approximate ratio of 1:4 every 3-4 days. Promega CellTiter-Glo 3D Cell Viability Assay was used to characterize the short-term cell viability in different formulation conditions. Cells were seeded at 10,000 cells per well in an opaque 96 well plate in 100 uL of media containing polymers of interest. MoNi_23%_, MPPhe_8%_ and MoPhe_6%_ polymers were added to media at concentrations of 10, 5, 2.5, 1, 0.5, and 0.1 mg/mL. Four wells were used as replicates for each polymer concentration. Relative viability was measured after 1 day in culture by adding 100 uL per well of the CellTiter-Glo reagent, mixing for 5 minutes, allowing the plate to sit for 25 minutes, and then reading the luminescent signal with a 1 second integration time. LC_50_ values were determined for each polymer by plotting the absolute signal for each of the four replicates in GraphPad Prism 9 and using the [Agonist] vs. Response – Find ECanything nonlinear fit. Fit parameter F was constrained to 50, Bottom was constrained to 4 (the negative control for the assay) and the Top was constrained to be the same for all data sets (cell viability should be equal for all data sets as polymer concentration approaches 0).

### Streptozotocin (STZ) induced model of diabetes in rats

Male Sprague Dawley rats (Charles River) were used for experiments. Animal studies were performed in accordance with the guidelines for the care and use of laboratory animals; all protocols were approved by the Stanford Institutional Animal Care and Use Committee. The protocol used for STZ induction adapted from the protocol by Kenneth K. Wu and Youming Huan,^20^ and have been previously described.^13, 21^ Detailed methods are included in the supplemental information. Diabetes was defined as having 3 consecutive blood glucose measurements >300 mg/dL in non-fasted rats.

### In vivo pharmacodynamics in diabetic rats

Diabetic rats were fasted for 4-6 hours. For initial blood glucose studies rats were injected subcutaneously (1.5U/kg) with the following formulations: (i) Humulin, or (ii) Humulin with 0.01 wt.%AC/DC excipient (MoNi_23%_, MpPhe_8%_, MoPhe_6%_). Humulin formulations were tested at 6 aging time points of 0, 0.5, 1, 2, 4, and 6 months, and Humulin with AC/DC excipient was tested at 0, 2, 4, and 6 months of aging. The preparation of formulations was staggered so that all samples reached their endpoint age at the same time and all aging timepoints could be compared in the same cohort of rats. 32 rats with fasting glucose levels >300 mg/dL were randomized to a formulation group (8 rats/group) and each rat received that formulation at all levels of aging (the order of the aging timepoints rats received was also randomized). For blood glucose studies after formulation aging at 50 °C, rats were injected subcutaneously (1.5U/kg) with the following formulations: (i) Humulin, (ii) Aged Humulin (t = 1 day), (iii) Humulin + MoNi_23%_, or (iv) Aged Humulin + MoNi_23%_ (t = 4 days). 16 Rats with fasting glucose levels >300 mg/dL were randomized to either the Humulin control group or the MoNi_23%_ group. Within both groups, the order that the aged formulations were given was also randomized and formulations were administered on separate experimental days. Before injection, baseline blood glucose was measured. After injection, blood was sampled every 30 minutes for 5 hours. Blood glucose was measured using a handheld blood glucose monitor. The maximum change in blood glucose measured from baseline was used as a metric of bioactivity of each formulation to assess *in vivo* bioactivity after aging.

### In vivo pharmacokinetics in diabetic rats

Diabetic rats were fasted for 4-6 hours. For pharmacokinetic studies rats were injected subcutaneously (1.5U/kg) with the following formulations: (i) Humulin, (ii) Aged Humulin (t = 6 months), (iii) Humulin + MoNi_23%_, or (iv) Aged Humulin + MoNi_23%_ (t = 6 months). 16 diabetic rats were randomized to a formulation group: Humulin or Humulin + MoNi_23%_ (8 rats/group). Within each group, rats received both the fresh (t = 0 months) or aged (t = 6 months) formulations in a randomized order. After subcutaneous injection, blood was sampled every 15 minutes for 2 hours and blood was collected in serum tubes (Sarstedt) for analysis with ELISA. Serum insulin concentrations were quantified using a Human Insulin ELISA kit (Mercodia).

### Statistics

All data is shown as mean ± standard error unless specified. For the *in vitro* activity assay (AKT) EC_50_ regression (log(agonist) vs. response (three parameters)) was plotted using GraphPad Prism 8. GraphPad Prism 8 Extra sum-of-squares F-test was used to test if Log(EC_50_) differed between datasets. Data sets were compared in pairs, and Bonferroni post-hoc tests were used to adjust for multiple comparisons (alpha=0.008). For blood glucose measurements, a REML repeated measures mixed model was used to test for differences at different aging timepoints within a formulation (JMP Pro 14). Rat was included as a random effect and the age of the formulation as a within-subject fixed effect. A post-hoc Tukey HSD test was used on Humulin formulations to determine statistical significance between aging timepoints.

## RESULTS AND DISCUSSION

### AC/DC excipient insulin stabilizing mechanism

We selected our three top performing candidates from a previous screen of amphiphilic AC/DC excipients for their ability to stabilize monomeric insulin.^19^ These excipients were composed of either acryloylmorpholine (Mo) or methoxypropylacrylamide (Mp) as a hydrophilic carrier monomer copolymerized with either N-Isopropylacrylamide (Ni) or phenylacrylamide (Phe) as a hydrophobic dopant monomer. This excipient design is hypothesized to preferentially occupy the air-water interface and consequently inhibit insulin-insulin interactions occurring at these interfaces (Figure 1). We sought to evaluate this hypothesis through time-resolved surface tension and interfacial rheology experiments with a model AC/DC excipient, poly(acryloylmorpholine_77%_-*co*-N-Isopropylacrylamide_23%_) (MoNi_23%_), co-formulated with commercial Humulin R (Eli Lilly) (Figure 2).

**Figure 1.**
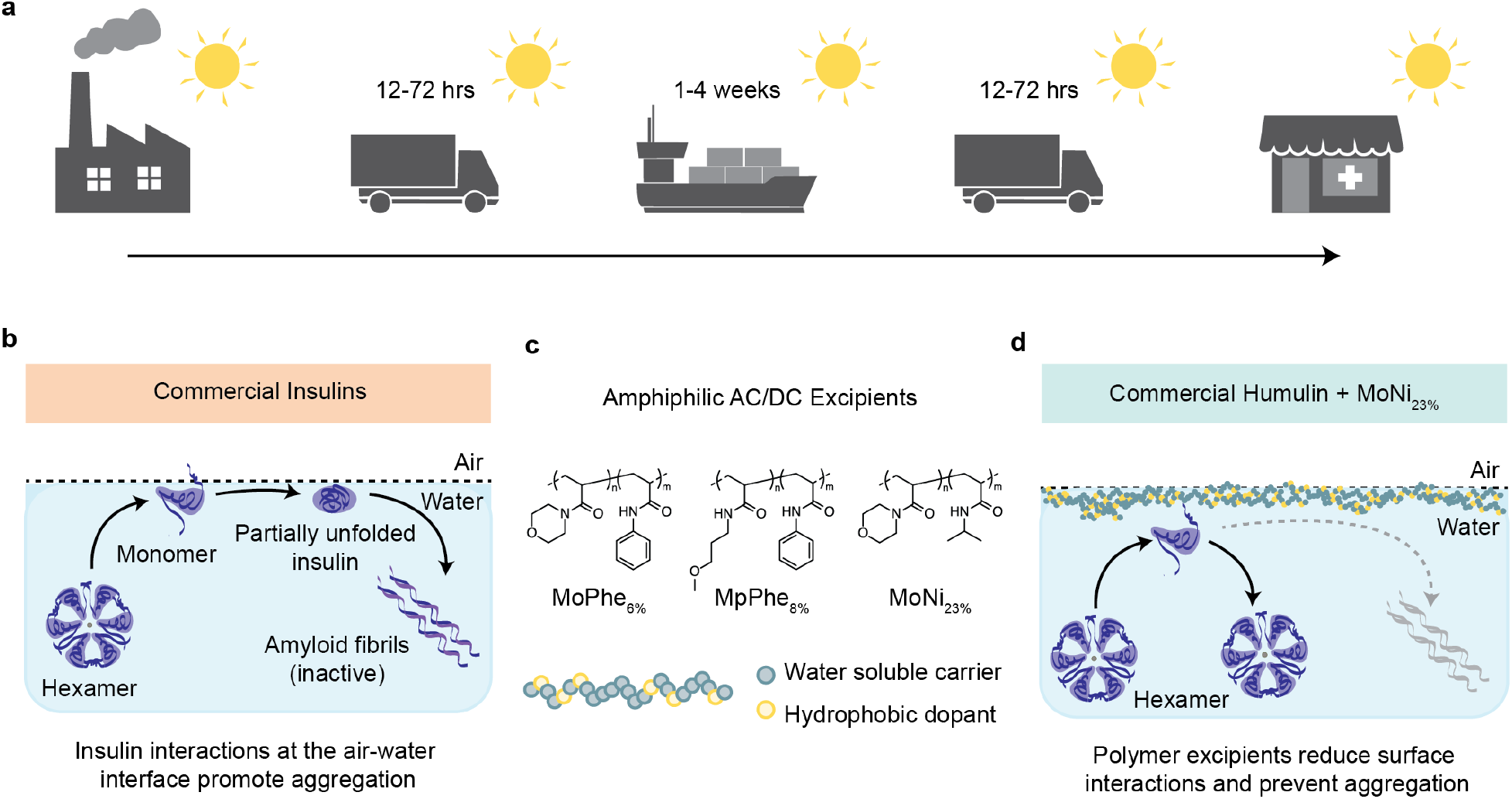
Scheme of the cold chain and insulin aggregation mechanism. **a,** To maintain integrity, commercial insulin formulations must currently be transported and stored in refrigerated containers for the weeks-long duration of worldwide distribution. **b,** Aggregation mechanism of commercial insulin formulations. The insulin hexamer is at equilibrium with monomers in formulation. These monomers interact at the interface, where the exposure of hydrophobic domains during insulin-insulin interaction nucleate amyloid fiber formation. **c,** Chemical structure of AC/DC excipients, poly(acryloylmorpholine_77%_-*co*-N-isopropylacrylamide_23%_) (MoNi_23%_). AC/DC excipients have a molecular weight between 2-5 kDa (See Table S1). **d,** AC/DC excipients are amphiphilic copolymers that adsorb to interfaces, reducing insulin-insulin interactions and delaying the nucleation of insulin amyloidosis.

**Figure 2:**
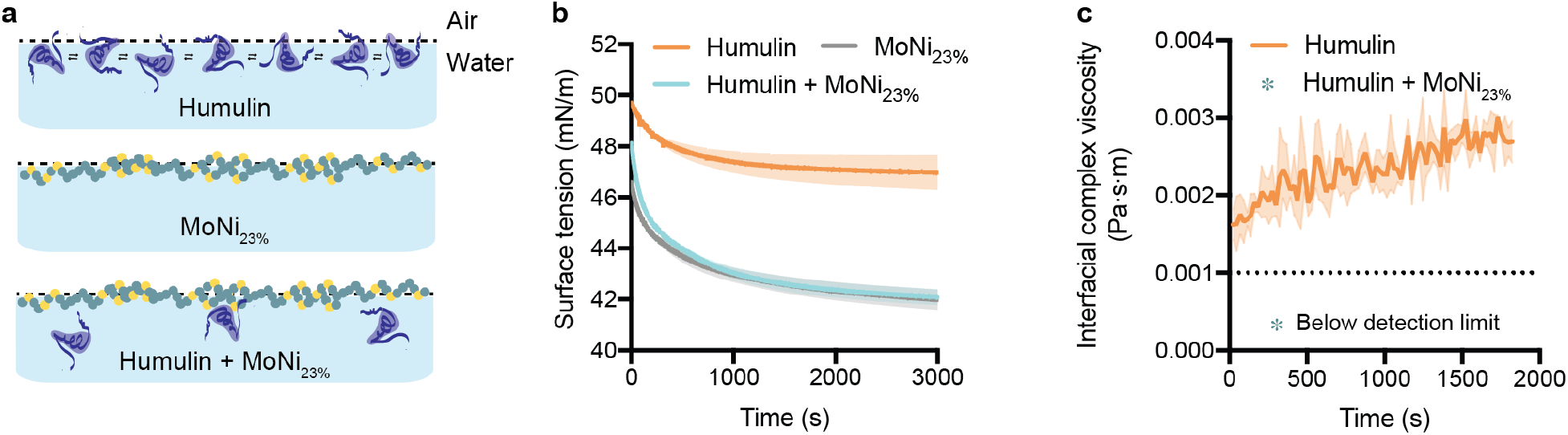
Experimental insight into the mechanism of AC/DC excipient stabilization. **a,** Illustration of proposed stabilization mechanism: (i) In commercial Humulin, monomers at the interface have associative interactions. (ii) Alone, MoNi_23%_ occupies the interface without the presence of insulin. (iii) In combination with Humulin formulations, MoNi_23%_ disrupts insulin-insulin surface interactions, providing a mechanism for inhibiting aggregation. **b,** Surface tension measurements of Humulin, MoNi_23%_ (0.01 wt.%) formulated with formulation excipients, and Humulin formulated with MoNi_23%_ (0.01 wt.%) (n=2). **c,** Interfacial rheology measurements of Humulin. Measurements for Humulin formulated with MoNi_23%_ (0.01 wt.%) fell below the resolution of the instrument, indicating that there is no protein aggregation at the interface (n=3).

Equilibrium surface tension measurements of Humulin R, Humulin R containing MoNi_23%_ (0.01 wt.%), and a solution of MoNi_23%_ (0.01 wt.%) containing the same formulation excipients (i.e., Humulin R without the insulin) revealed that the presence of MoNi_23%_ resulted in surface tension values well below Humulin R (approximately 42 vs. 47 mN/m, Figure 2b). Moreover, a ten-fold increase in the MoNi_23%_ concentration (0.1 wt.%) further reduced the surface tension of the formulation (Supplementary Figure 1A). The decrease in surface tension upon addition of MoNi_23%_ to Humulin indicates that there are more species at the interface when MoNi_23%_ and Humulin are formulated together, compared to Humulin alone. The decreased surface tension concomitant with the increased concentration of MoNi_23%_ in the absence of Humulin indicates that the surface is not saturated at 0.01 wt.% MoNi_23%_. However, the surface tension is identical for formulations of Humulin and MoNi_23%_ and MoNi_23%_ with formulation excipients, indicating that there are similar number of molecular species at the interface regardless of the presence of Humulin. Further, the addition of completely hydrophilic poly(acryloylmorpholine) (Mo) to Humulin did not lower the surface tension, indicating that the amphiphilic copolymer is required to displace insulin (Supplementary Figure 1B). Together, these surface tension experiments help demonstrate that MoNi_23%_ preferentially adsorbs and dominates the air-water interface.^22^

Interfacial shear rheology measurements demonstrated that addition of MoNi_23%_ (0.01 wt.%) to Humulin R reduced interfacial complex viscosity to below the detection limit of the instrument compared to Humulin R, which exhibited values between 0.002 - 0.003 Pa·s·m (Figure 2c). The complex viscosity of Humulin is indicative of associative insulin-insulin interactions that can dissipate viscous energy at the interface. While not quantitative, the lowering of the interfacial complex viscosity below instrument detection limits is indicative that the addition of MoNi_23%_ disrupts insulin-insulin interactions at the interface.

When this complex interface is subjected to interfacial stresses and agitation, it is likely that these insulin-insulin associations can nucleate amyloid fibril formation and lead to aggregation. Together, the surface tension and interfacial rheology experiments suggest a mechanism of AC/DC enhanced insulin stabilization where preferential adsorption of the AC/DC excipient to the air-water interface disrupts insulin-insulin interactions.

### AC/DC excipients for long-term stability of insulin

We then sought to evaluate the capacity of our three previous top performing AC/DC excipients, MoNi_23%_, poly(acryloylmorpholine_94%_-*co*-phenylacrylamide_6%_) (MoPhe_6%_) and poly(methoxypropylacrylamide_92%_-*co*-phenylacrylamide_8%_) (MpPhe_8%_) to act as simple “drop-in” excipients to stabilize Humulin R through stressed aging. Formulations of Humulin alone or Humulin with an AC/DC excipient added were prepared and aged for 0, 2, 4, or 6 months at 37 °C with constant agitation (150 rpm on an orbital shaker plate). The preparation of formulations was staggered so that all samples reached their endpoint age at the same time. Both visual inspection and a transmittance assays were used to determine if the insulin had aggregated (Figure 3). ^11, 13^ Insulin aggregates scatter light, and thus aggregation can be defined as a change in transmittance greater than 10%.^11, 13^ Humulin alone began to aggregate after 2 weeks of stressed aging. In contrast, all insulin formulations containing AC/DC excipients MoPhe_6%_, MpPhe_8%_, and MoNi_23%_ at concentrations of 0.01, 0.1 or 0.5 wt.% did not show any signs of insulin aggregation over the course of the 6-month study, with the exception of MpPhe_8%_ at 0.5 wt.% (Figure 3b, Supplementary Figure 2). Thus, to minimize the amount of copolymer excipient in formulation, only the 0.01 wt.% formulations were used for the rest of the studies reported here.

**Figure 3:**
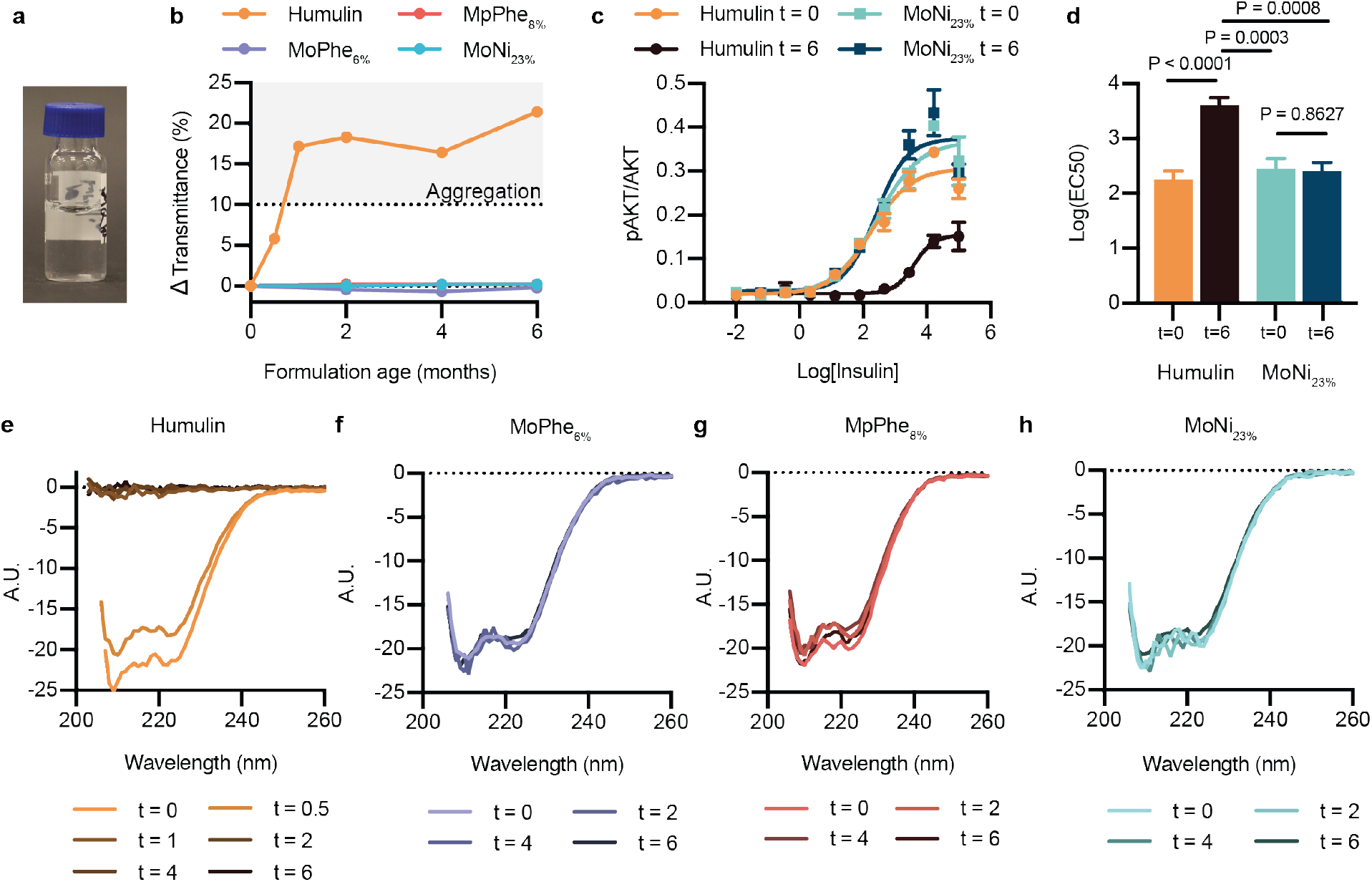
Formulation with AC/DC copolymers stabilizes insulin. **a,** 1mL of Commercial Humulin or Humulin with the addition of AC/DC excipients (i) MoPhe_6%_, (ii) MpPhe_8%_, (iii) MoNi_23%_ were aliquoted into 2mL glass vials and aged at 37°C with constant agitation (150 rpm) for 0, 2, 4, and 6 months. Additional 2 week and 1 month timepoints were added for the Humulin control. All formulations were at a concentration of 95U/mL (diluted so that copolymers could be added to commercial Humulin). **b,** Transmittance assay to assess the aggregation of proteins in formulation over time by monitoring changes in transmittance at 540nm (n=1 per formulation timepoint). **c,** *In vitro* activity by assaying for phosphorylation of Ser^473^ on AKT after stimulation with either Humulin or MoNi_23%_ at 0 month and 6 month timepoints. Insulin concentrations are shown as Log(ng/mL). **d,** Log(EC_50_) values for each formulation. Statistical significance was assessed using the Extra sum-of-squares F-test to determine if Log(EC_50_) differed between datasets. Data sets were compared in pairs, and Bonferroni post-hoc tests were used to adjust for multiple comparisons (alpha=0.008). **e-h,** Circular dichroism spectra from 200-260nm for each formulation (diluted to 0.2 mg/mL in PBS) at each time point. **c,** Results shown are mean ± s.e plotted as a ratio of [pAKT]/[AKT] for each sample (n=3 cellular replicates) and an EC_50_ regression (log(agonist) vs. response (three parameters)) was plotted using GraphPad Prism 8. **d,** Statistical significance was assessed using the Extra sum-of-squares F-test to determine if Log(EC_50_) differed between datasets. Data sets were compared in pairs, and Bonferroni post-hoc tests were used to adjust for multiple comparisons (alpha=0.008).

To further validate the transmittance results, which only assess insulin aggregation, we evaluated *in vitro* activity by assaying for phosphorylation of Ser^473^ on protein kinase B (AKT) after stimulating C2C12 cells with either Humulin or Humulin containing MoNi_23%_ (0.01 wt.%) at both the 0 month and 6 month timepoints (Figure 3c-d). Fresh formulations and the aged Humulin+MoNi_23%_ formulation showed equivalent bioactivity (Humulin_t=0_ Log(EC_50_) = 2.252 ± 0.158; MoNi_23%,t=0_ Log(EC_50_) = 2.448 ± 0.186; MoNi_23%,t=6_ Log(EC_50_) = 2.405 ± 0.158), whereas aged Humulin R exhibited almost complete loss of bioactivity (Humulin_t=6_ Log(EC_50_) = 3.606 ± 0.139) (Figure 3c-d).

While these *in vitro* AKT assay results supported the transmittance data, we sought to further confirm insulin formulation integrity by using circular dichroism to observe insulin secondary structure for each formulation timepoint (Figure 3e-h). Formulations stabilized with the AC/DC excipients exhibited no changes in secondary structure after stressed aging, whereas Humulin alone had lost all structural features by 1 month. These data corroborate both the transmittance and *in vitro* activity data.

### Bioactivity of aged insulin in diabetic rats

To evaluate the integrity of the aged insulin formulations in a functional setting *in vivo*, we assessed formulation activity in diabetic rats. Administration of streptozotocin was used to induce insulin-dependent diabetes in a cohort of 32 male rats. These rats were randomly assigned to one of four formulation groups: (i) Humulin, or Humulin comprising either (ii) MoPhe_6%_, (iii) MpPhe_8%_, or (iv) MoNi_23%_ at 0.01 wt.%, and each rat received that formulation at each aging timepoint (0, 2, 4, 6 months). The preparation of formulations was staggered so that all samples reached their endpoint age at the same time and all aging timepoints could be compared in the same cohort of rats. Insulin was administered subcutaneously in fasted rats (1.5U/kg) and blood glucose levels were measured every 30 minutes. Active formulations resulted in a distinct initial drop in blood glucose from extreme hyperglycemia that reached a minimum in the range of normoglycemia between 60-100 minutes after administration (Figure 4, Supplementary Figure 3). After this phase, blood glucose levels began to rise as insulin was cleared. In contrast, formulations that appeared aggregated in the *in vitro* transmittance assays following aging did not show this distinct reduction in glucose reminiscent of insulin action and instead resulted in a gradual decrease in glucose levels. The gradual decrease in glucose may suggest that some of the insulin is initially trapped in reversible aggregates, and over time these aggregates dissociate and result in a slow-acting insulin effect. The maximum difference in blood glucose from baseline to the minimum glucose levels was plotted for each formulation as a measure of formulation potency. All copolymer-stabilized formulations showed no difference in activity between aging timepoints, but Humulin alone demonstrated a large difference between aging timepoints (F_3,21_=23.83, P<0.0001), where a post-hoc Tukey HSD test revealed that Humulin after 2, 4, and 6 months of aging had decreased activity compared to fresh Humulin (t=0 months). These observations were corroborated by evaluation of insulin pharmacokinetics, where no differences were observed between fresh Humulin R (t=0 months) and Humulin+MoNi_23%_ initially (t=0 months) and after 6 months of aging, but a decrease in exposure was observed for the aged Humulin (t=6 months) (Figure 4f, Supplementary Figure 4). These data suggest that AC/DC excipients function as stabilizing ingredients for commercial formulations such as Humulin R without altering the insulin pharmacokinetics or pharmacodynamics.

**Figure 4:**
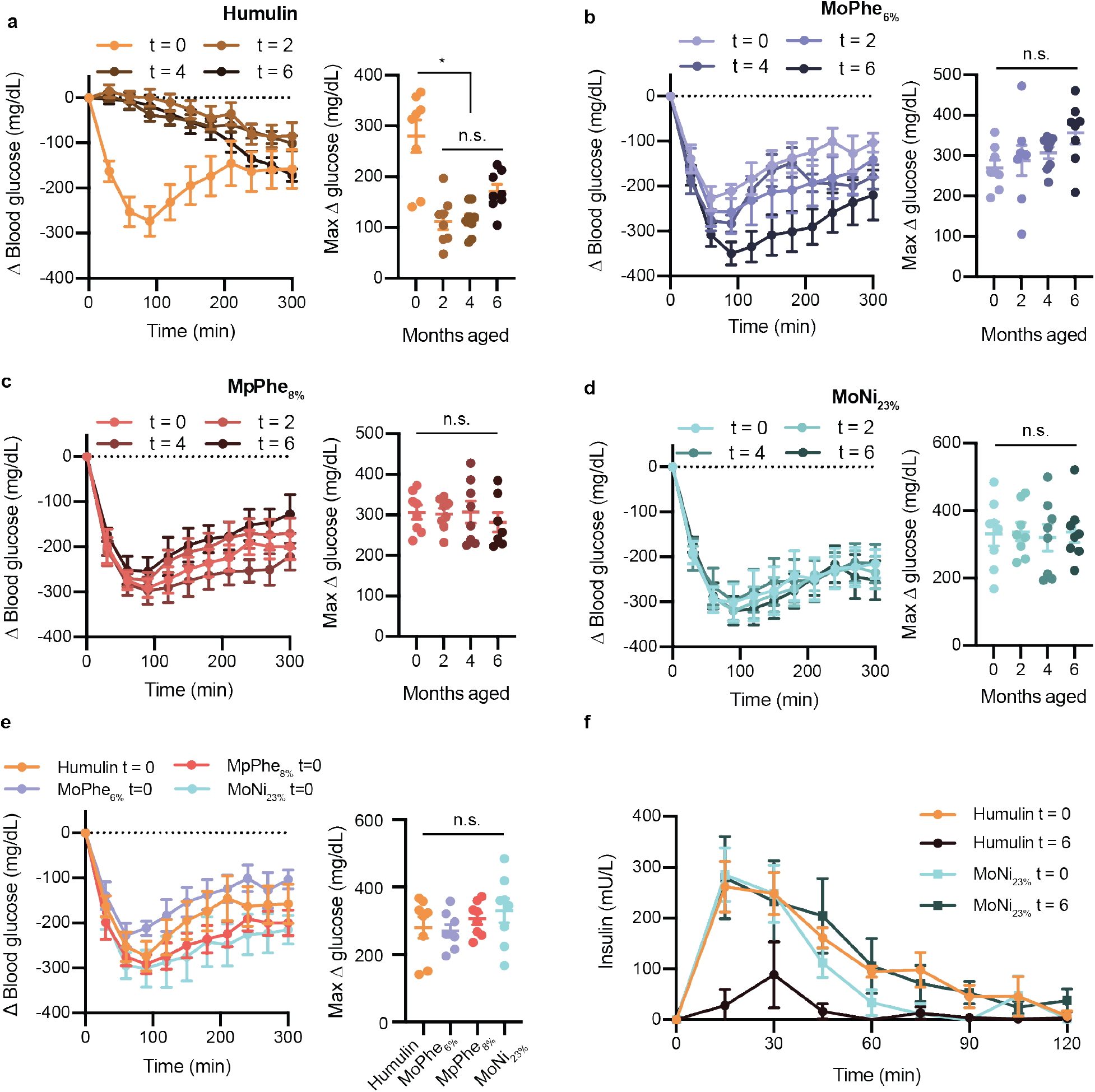
Insulin activity after aging in diabetic rats. Fasted diabetic male rats received subcutaneous administration (1.5U/kg) of each insulin formulation **a,** Humulin **b,** Humulin with MoPhe_6%_ **c,** Humulin with MpPhe_8%_, or **d,** Humulin with MoNi_23%_ at each aging timepoint (0, 2, 4, 6 months). **e,** Comparison of each formulation a t=0 months. In these assays, 32 rats were randomly assigned to one of the four formulation groups (n=8) and each rat received one dose of the formulation at each aging timepoint in a random order. Blood glucose levels were measured every 30 minutes using a handheld glucose monitor and the change in blood glucose relative to baseline glucose measurements were plotted. Baseline glucose measurements ranged from 300-600 mg/dL (See Supplementary Figure 5 for raw glucose curves). The maximum difference in glucose from baseline (Δ glucose) was also plotted for each formulation as a measure of formulation potency. **f,** Pharmacokinetics of Humulin and Humulin with MoNi_23%_ at t=0 and t=6 months. All data is shown as mean ± s.e. Statistical significance between max Δ glucose was assessed using a REML repeated measures mixed model with rat as a random effect and the age of the formulation as a within-subject fixed effect. A post-hoc Tukey HSD test was used on Humulin formulations to determine statistical significance between aging timepoints.

### High temperature aging of insulin formulations

To determine the capacity of AC/DC excipients to improve insulin cold-chain resilience, we evaluated the extent of stability imbued by one of our excipients, MoNi_23%_, under extreme manufacturing and distribution conditions (37 °C and 50 °C with constant agitation) (Figure 5). MoNi_23%_ was selected for further testing following cytotoxicity tests that revealed MoNi_23%_ and MoPhe_6%_ had more favorable LC_50_ values compared to MPPhe_8%_ (Supplementary Figure 6). These results, combined with previous success using MoNi_23%_ to stabilize monomeric insulin, ^19^ led us to choose to advance MoNi_23%_ through the extreme aging tests. Temperatures were selected to represent the temperature on a hot summer day (37 °C), and the upper temperature range that a shipping container or truck without refrigeration or insulation could reach during the peak of summer (50 °C).^23^ Humulin R can be purchased in 10mL glass vials that are packaged and shipped in cardboard boxes (Figure 5a, Supplementary Figure 7). MoNi_23_ (0.01 wt.%) was added to new vials of Humulin R using a syringe (dilution from 100 U/mL to 95U/mL to allow addition of copolymer; control vial was diluted with water) and the vials were then replaced in the original cardboard packaging with the package insert (Supplementary Figure 7a). The cardboard packaging was affixed to a rotary shaker inside a temperature-controlled incubator and agitated at 150 RPM (Supplementary Figure 7b).

**Figure 5:**
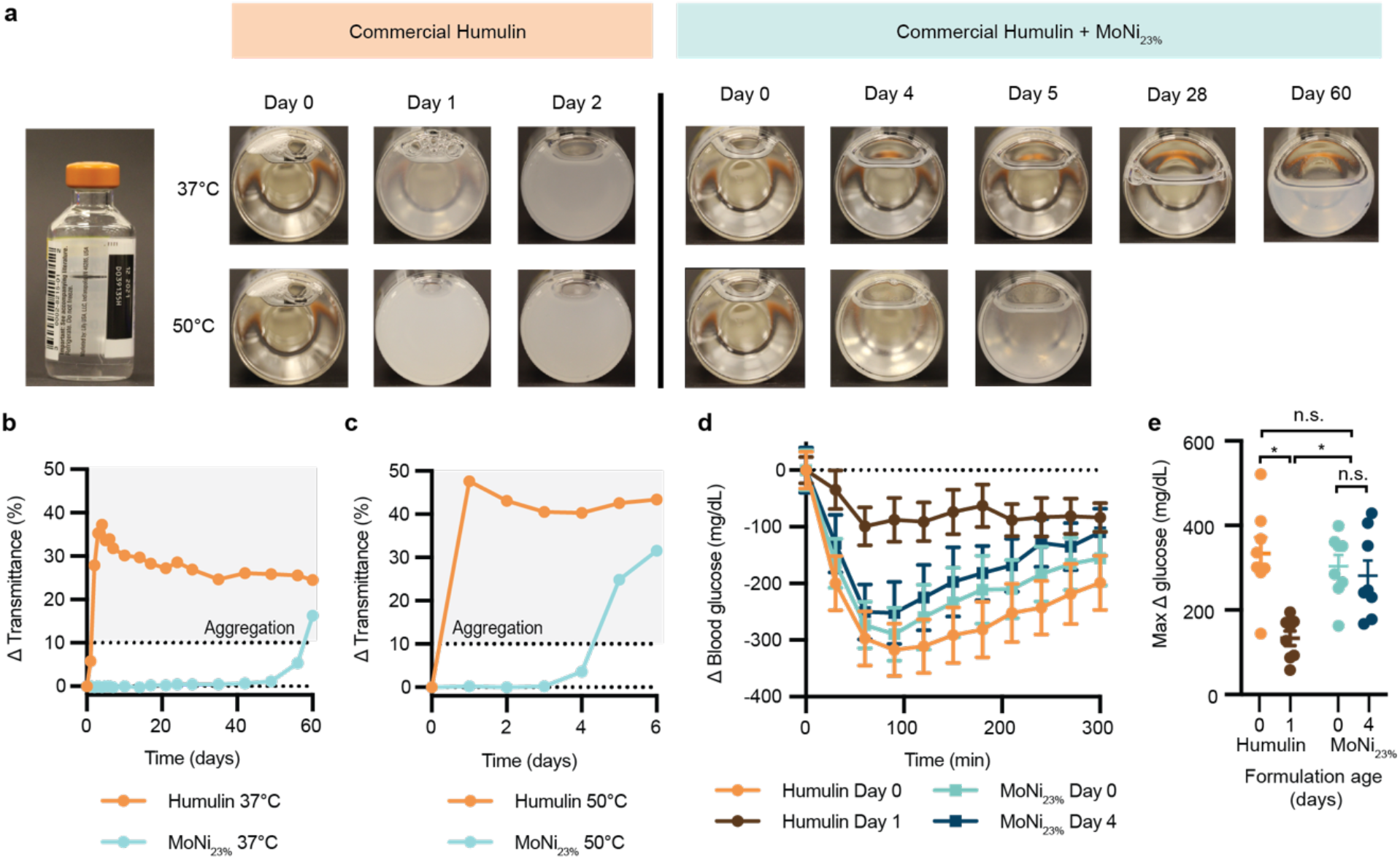
Stressed aging in commercial packaging. **a,** Humulin is often sold in standardized 10mL glass vials and packaged in cardboard boxes. Here we tested the stabilizing capacity of AC/DC copolymers in commercial packaging conditions under stressed conditions. 10mL vials of U100 Humulin R were diluted to 95 U/mL with the addition of 50 uL of a MoNi_23%_ stock solution (to a final concentration of 0.01 wt.% copolymer) or water (control). Dilution was necessary to add copolymer to the formulation. These vials were replaced in their original boxes and taped to a shaker plate (150 rpm) in a 37 °C (n=1 per formulation) or 50 °C (n=1 per formulation) incubator. Samples were observed and imaged daily. **b,c,** Transmittance assays for Humulin or Humulin comprising MoNi_23%_ after aging at **(b)** 37 °C and **(c)** 50 °C. Single samples (n=1) were tested for each transmittance curve. **d,** Blood glucose curves and **e,** maximum change in blood glucose (Δ glucose) in fasted diabetic rats for samples aged at 50 °C. Data is shown as mean ± s.e. Statistical significance between max Δ glucose was assessed using a REML repeated measures mixed model with rat as a random effect and the age of the formulation as a within-subject fixed effect. A post-hoc Tukey HSD test was used to determine statistical significance between aging timepoints and groups.

Visual inspection combined with a transmittance assay were used as the primary measures of insulin integrity (Figure 5a). These assays are consistent with our earlier experiments that demonstrated that the transmittance readings correlate well with both *in vitro* and *in vivo* functional activity assays. At 37 °C, Humulin alone begins to show visual changes in opacity at day 1 and becomes fully opaque by day 2. In contrast, when formulated with MoNi_23%_, the insulin formulation shows no visual changes in opacity until day 56 and remains below a 10% change in transmittance for 56 days. At 50 °C, commercial Humulin became fully opaque within one day. In contrast, formulation with MoNi_23%_ extends stability under these extreme conditions to past 4 days before the formulation becomes cloudy on day 5. These qualitative observations were consistent with quantitative transmittance readings (Figure 5b-c).

To verify functional insulin activity after aging at 50 °C *in vivo*, these formulations were evaluated in diabetic rats. The ability of (i) Humulin (t = 0 day), (ii) aged Humulin (t = 1 day), (iii) Humulin with MoNi_23%_ (t = 0 day), or (iv) aged Humulin with MoNi_23%_ (t = 4 days) to decrease glucose levels was measured in fasted diabetic rats. After subcutaneous administration of formulations (1.5U/kg), blood glucose levels were measured every 30 minutes (Figure 5d). Humulin (t = 0 day), Humulin with MoNi_23%_ (t = 0 day), and aged Humulin with MoNi_23%_ (t = 4 day) demonstrated an initial blood glucose drop that reached a minimum between 60-100 minutes after injection. These results were consistent with active formulations in earlier experiments. This characteristic glucose drop was absent in rats who received aged Humulin (t = 1 day), consistent with inactive formulations in earlier experiments. The maximum difference in glucose from baseline was also plotted for each formulation as a metric of formulation potency (Figure 5e).

Statistical analysis identified a difference between the potency of these formulations (F_3.18.18_=10.71, P=0.0003), whereby a post-hoc Tukey HSD test revealed that aged Humulin alone had significantly decreased activity compared to the other formulations. In contrast, there was no statistical difference between unaged Humulin, unaged Humulin with MoNi_23%_, as well as aged Humulin with MoNi_23%_ after stressed aging at 50 °C for 4 days.

## CONCLUSIONS

In this study, we report on the application of AC/DC excipients to imbue long term stability and cold chain resilience to commercial insulin formulations. While commercial insulin formulations have good shelf lives when stored properly, interruptions in the cold chain can decrease insulin bioactivity and formulation integrity. The primary mechanism of insulin destabilization is through insulin adsorption at the air water interface, where insulin-insulin interactions promote amyloid fibril nucleation.^6–7, 9^ We hypothesized that amphiphilic copolymer excipients could potentially displace insulin from the interfacial region, and thus improve stability by reducing the probability of nucleation of insulin-insulin aggregation events at the interface. Surface tension and interfacial rheology data on our top-performing excipient MoNi_23%_ support this hypothesis by showing that these copolymers preferentially adsorb to the air-water interface and disrupt insulin-insulin interactions. No thermo-responsive behavior was observed by the MoNi_23%_ excipient at the temperatures tested in this study (Figure S8), thus disruption of surface interactions remains the most likely mechanism for stability.

When used as simple, low concentration (0.01 wt.%) formulation additives, AC/DC excipients preserve insulin activity through 6 months of stressed aging without modifying formulation pharmacokinetics, protein secondary structure, formulation clarity, or *in vivo* bioactivity. When further subjected to harsh stressed aging tests in standard packaging, Humulin R formulated with our MoNi_23%_ excipient did not aggregate for over 56 days of constant agitation at 37 °C, and four days of constant agitation at 50 °C, whereas Humulin R alone aggregated within two days at 37 °C and within one day at 50 °C. While agitation experienced by pharmaceuticals during typical storage and transportation can be highly complex, these processes can be mimicked in vitro using elevated temperatures and continuous agitation. Stressed aging, through incubation at elevated temperatures with continuous agitation, has been previously used to test insulin stability.^11, 13–14, 19, 21^ Even in hot climates with limited cold chain infrastructure, it is unlikely shipping containers would remain at 50 °C with continuous agitation with no reprieve for over four days and nights. Further, in our harsh stressed aging studies we use 10 mL vials, which resulted in a higher rate of interfacial exchange compared to initial stability studies in autosampler vials due to a larger diameter. The increased stability observed for Humulin R formulated with our MoNi_23%_ excipient compared to Humulin R alone suggests that our MoNi_23%_ excipient has utility in stabilizing insulin under various agitation conditions where interfacial turnover may be higher (i.e., horizontal agitation). The conditions evaluated in this study represent extreme exposure conditions during shipping in uninsulated containers or trucks in the hottest climates in the world where transport can take weeks before reaching patients.^18^ In addition to refrigerated transport in the early stages of the cold chain, maintaining proper transport and storage conditions during local distribution and once in patients’ hands presents a challenge in many parts of the world.^24^ This work suggest that the addition of AC/DC excipients can preserve insulin formulation integrity during even severe cold chain interruptions, enabling a reduction in cold chain requirements for insulin transportation and storage that are difficult to maintain in under-resourced environments.^3, 24–29^

These AC/DC excipients are synthesized at molecular weights below the glomerular filtration threshold to ensure they are eliminated without tissue accumulation, and they are prepared with facile and scalable controlled radical polymerization techniques from inexpensive starting materials. Previous exploration of this class of amphiphilic copolymer showed that they do not alter blood chemistry markers for liver and kidney function following repeated administration in rats, and exhibit cytotoxicity levels comparable to the phenolic preservatives used in commercial insulin formulations.^19^

Taken together, the results reported herein demonstrate the potential of AC/DC excipients as insulin formulation additives to improve cold chain resilience, thereby expanding global access to these critical drugs for treatment of diabetes. Further, based on the identified stabilization mechanism, AC/DC excipients are promising candidates to stabilize other biopharmaceuticals, such as monoclonal antibodies, that lose bioactivity as a result of aggregation at interfaces.^22, 30^ Future studies will require continued evaluation of the limits of AC/DC stabilized insulin and explore the application of these excipients to other protein therapeutics.

## Supporting Information

Supporting information document contains the following:

- Supplemental Methods
- Figure S1
- Figure S2
- Figure S3
- Figure S4
- Figure S5
- Figure S6
- Figure S7
- Figure S8
- Table S1

## AUTHOR INFORMATION

### Corresponding Author

* Person to whom correspondence should be addressed: Dr. Eric A. Appel

### Present Addresses

† Present Address for A.A.A.S: Department of Health Technology, Technical University of Denmark, 2800 Kgs. Lyngby, Denmark

### Author Contributions

‡ C.L.M and J.L.M contributed equally. Conceptualization, C.L.M., J.L.M., E.A.A. Methodology, C.L.M., J.L.M., A.K., C.M.M., A.A.A.S. Investigation, C.L.M., J.L.M., A.K., C.M.M., B.S.O. Writing – Original Draft, C.L.M and J.L.M. Writing – Review & Editing, C.L.M., J.L.M., A.K., C.M.M., B.S.O., A.A.A.S., G.G.F., E.A.A. Supervision E.A.A.

## ACKNOWLEDGMENT

The authors thank the Veterinary Service Centre staff for their technical assistance. The authors also acknowledge the High Throughput Bioscience Center (HTBC) at Stanford Medicine.

## For Table of Contents Use Only

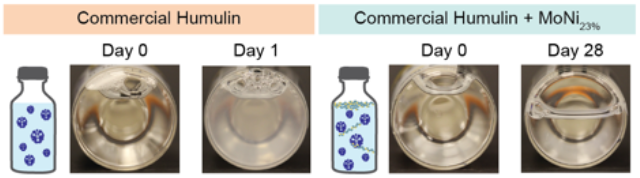

