## Supplemental Information for "Engineering Insulin Cold Chain Resilience to Improve Global Access"

### SUPPLEMENTARY METHODS

#### *Polymer Synthesis*

The procedure to synthesize $MoNi_{23\%}$ AC/DC excipient is as follows and is nearly identical for all other carrier/dopant combinations, where only the carrier/dopant selection and concentrations are changed. MORPH (566 mg, 4.02 mmol, 36.5 eq.), NIPAM (168 mg, 1.485 mmol, 13.5 eq.), 2CPDT (38 mg, 0.11 mmol, 1 eq.) and AIBN (3.6 mg, 0.02 mmol, 0.2 eq.) were combined and diluted with DMF to a total volume of 2.25 mL (33.3 w/v vinyl monomer concentration) in an 8 mL scintillation vial equipped with a PTFE septa. The reaction mixture was sparged with nitrogen gas for 10 minutes and then heated for 12 hours at 65 °C. To remove the Z-terminus of the resulting copolymer, AIBN (360 mg, 2.2 mmol, 20 eq.) and LPO (88 mg, 0.22 mmol, 2 eq.) were added to the reaction mixture, which was then sparged with nitrogen gas for 10 minutes and heated for 12 hours at 90 °C (*26*). Z-group removal was confirmed by the ratio of the refractive index to UV (λ = 310 nm) intensity in size exclusion chromatography (SEC) analysis. Resulting copolymers were precipitated three times from ether and dried under vacuum overnight.

#### *In vitro insulin cellular activity assay:*

In vitro insulin activity was tested using the AKT phosphorylation pathway using AlphaLISA SureFire Ultra (Perkin-Elmer) kits for detection of phosphorylated AKT 1/2/3 (pS473) compared to total Akt1. Humulin, Aged Humulin (t = 6 months) , Humulin + $MoNi_{23\%}$, and Aged Humulin + $MoNi_{23\%}$ (t = 6 months) formulations were tested. C2C12 mouse muscle myoblasts (ATCC CRL-1772) were cultured and were confirmed to be mycoplasma free prior to use. Dulbecco's Modified Eagle's Medium (DMEM) (Gibco; 4.5

g/L D-glucose, L-glutamine, 110 mg/L sodium pyruvate) was supplemented with 10% fetal bovine serum (FBS) and 5% penicillin-streptomycin. Cells were grown in a 96-well tissue culture plate for 24 hours (Seeding density= 25,000 cells/well in 200 μL culture media). Prior to insulin stimulation, the cells were washed twice with 200 μL of unsupplemented DMEM and starved in 100 μL of unsupplemented DMEM overnight. The media was then removed and the cells were stimulated with 100 μL of insulin (i) Humulin, (ii) Aged Humulin (t = 6 months) , (iii) Humulin + $MoNi_{23\%}$, or (iv) Aged Humulin + $MoNi_{23\%}$ (t = 6 months), diluted in unsupplemented DMEM, for 30 min while incubating at 37 °C. Cells were washed twice with 100 μL of cold 1X Tris-buffered saline before adding 100 μL of lysis buffer to each well and shaking for at least 10 minutes at room temperature to fully lyse cells. 30 μL of lysate was transferred to a 96-well white half-area plate for each assay. Assays were completed according to the manufacturer's protocol. Plates were incubated at room temperature and read 18-20 hours after the addition of the final assay reagents using a Tecan Infinite M1000 PRO plate reader. Results were plotted as a ratio of [pAKT]/[AKT] for each sample (n=3 cellular replicates) and an EC50 regression (log(agonist) vs. response (three parameters)) was plotted using GraphPad Prism 8.

***Ethical approval of studies including animal experiments***

All animal studies were performed in accordance with the guidelines for the care and use of laboratory animals; all protocols (Protocol No. 32873) were approved by the Stanford Institutional Animal Care and Use Committee prior to the research being conducted.

***Streptozotocin (STZ) induced model of diabetes in rats***

Male Sprague Dawley rats 180-250g (8-10 weeks) were weighed and fasted the morning of treatment (6-8 hours) prior to treatment with STZ in the afternoon. Pre-weighed STZ was protected from light and diluted to 10-20 mg/mL in 1 mL sodium citrate buffer (pH=4.5) immediately before injection. Rats were injected with STZ solution (65 mg/kg) intraperitoneally. Rats were given water containing 10% sucrose for 24 hours after administration of STZ. Three days after treatment with STZ, rat blood glucose levels were tested for hyperglycemia via tail vein blood collection using a handheld Bayer Contour Next glucose monitor (Bayer). Subsequent glucose monitoring was performed daily. Diabetes was defined as having 3 consecutive blood glucose measurements >300 mg/dL in non-fasted rats.

### SUPPLEMENTARY FIGURES

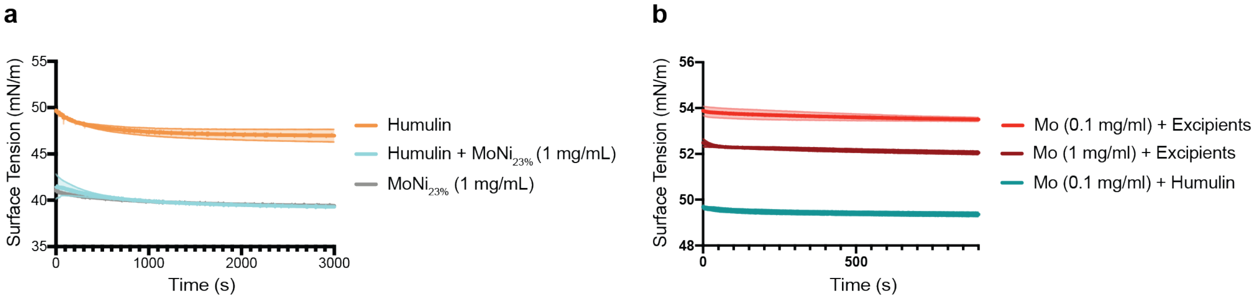


**Figure S1. Surface Tension polymer formulations. (A)** Surface tension measurements of Humulin (95U), $MoNi_{23\%}$ (0.1 wt.%) formulated with glycerol (1.6 wt.%) and metacresol (0.25 wt.%), and Humulin (95U) formulated with $MoNi_{23\%}$ (0.1 wt.%). **(B)** Surface tension measurements of hydrophilic poly(acryloylmorpholine) (Mo) at varying concentrations formulated with glycerol (1.6 wt.%) and metacresol (0.25 wt.%) (denoted "+ Excipients") or Humulin (95U).

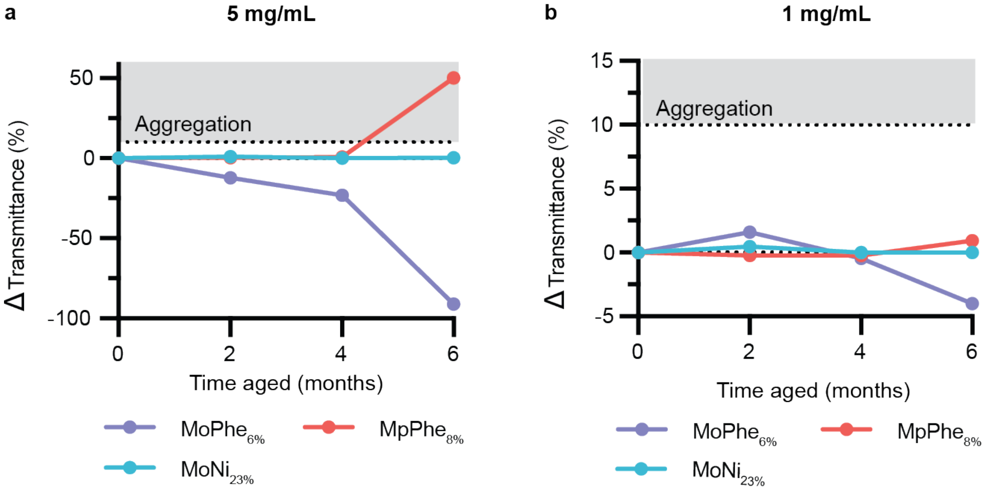


**Figure S2. Transmittance assays for 5 mg/mL and 1 mg/mL formulations.** 1mL of Humulin with the addition of AC/DC excipients (i) $MoPhe_{6\%}$, (ii) $MpPhe_{8\%}$, (iii) $MoNi_{23\%}$ were aliquoted into 2mL glass vials and aged at 37°C with constant agitation (150 rpm) for 0, 2, 4, and 6 months. **(A)** 5mg/mL polymer excipient in formulation (0.5 wt.%) and **(B)** 1 mg/mL polymer excipient in formulation (0.1 wt/%).

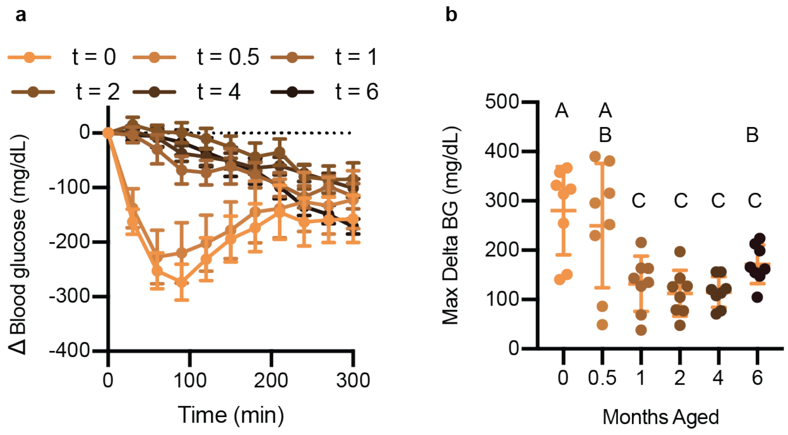


**Figure S3. Blood glucose curve for Humulin.** Humulin glucose curve including t=0.5 and t=1 month time points. Fasted diabetic male rats (n=8) received subcutaneous administration (1.5U/kg) of Humulin. 8 rats were randomly assigned to the Humulin group. Each rat received one dose of the formulation at each aging timepoint in a random order. Blood glucose levels were measured every 30 minutes using a handheld glucose monitor and the change in blood glucose relative to baseline glucose measurements was plotted. Statistical significance between max Δglucose was assessed using a REML repeated measures mixed model with rat as a random effect and the age of the formulation as a within-subject fixed effect. A post-hoc Tukey HSD test was used on formulations to determine statistical significance between aging timepoints. Groups not connected by the same letter are significantly different.

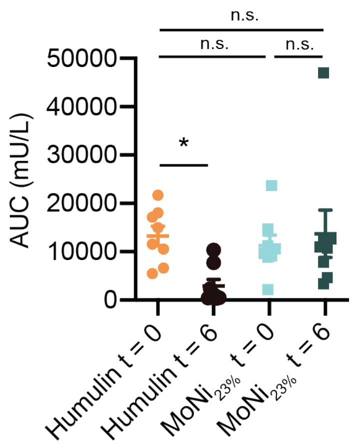


**Figure S4. Area under the curve for pharmacokinetics.** Area under the curve was calculated for the pharmacokinetic curves of Humulin, aged Humulin (t=6), Humulin + $MoNi_{23\%}$ and aged $MoNi_{23\%}$ (t=6). No difference was observed between fresh Humulin or polymer stabilized formulations. Aged Humulin showed decreased area under the curve compared to Humulin. All data is shown as mean ± s.e. Statistical significance between AUC was assessed using a REML repeated measures mixed model with rat as a random effect and the age of the formulation as a within-subject fixed effect. A post-hoc Tukey HSD test was used on formulations to determine statistical significance between aging timepoints.

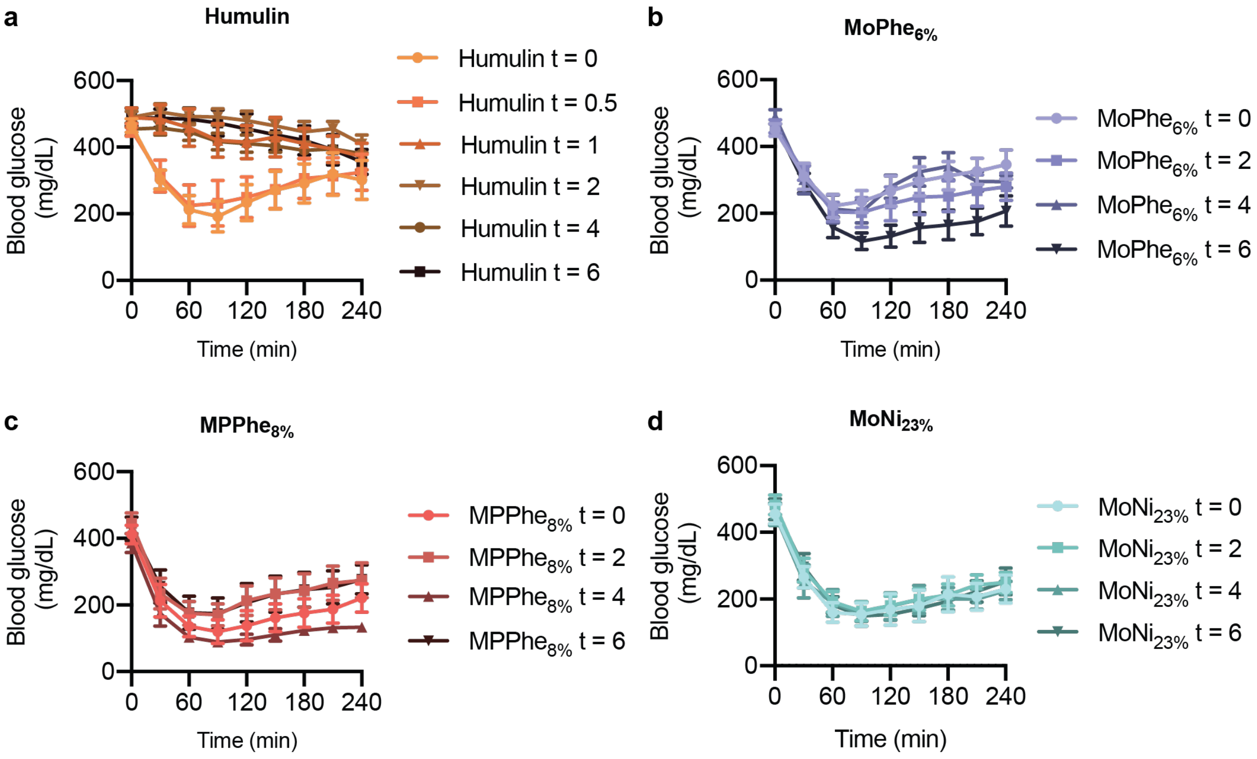


**Figure S5. Insulin activity after aging in diabetic rats.** Fasted diabetic male rats received subcutaneous administration (1.5U/kg) of each insulin formulation **a,** Humulin **b,** Humulin with $MoPhe_{6\%}$, **c,** Humulin with $MpPhe_{8\%}$, or **d,** Humulin with $MoNi_{23\%}$ at each aging timepoint. In these assays, 32 rats were randomly assigned to one of the four formulation groups (n=8) and each rat received one dose of the formulation at each aging timepoint in a random order. Blood glucose levels were measured every 30 minutes using a handheld glucose monitor. This data is presented as change from baseline glucose in Figure 4 of the main text.

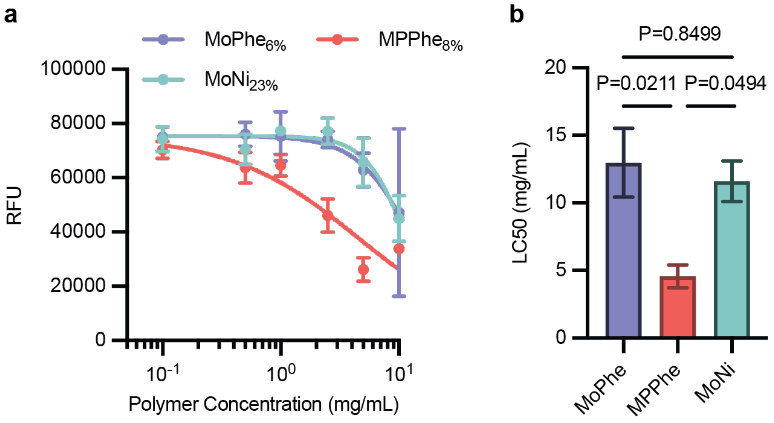


**Figure S6. Cytotoxicity and LC50 for AC/DC excipients. a,** NIH 3T3 cells were cultured with AC/DC polymers in cell media for 1 day and then cell viability assay was performed. $MoNi_{23\%}$, $MPPhe_{8\%}$ and $MoPhe_{6\%}$ polymers were added to media at concentrations of 10, 5, 2.5, 1, 0.5, and 1 mg/mL. Four wells were used as replicates for each polymer concentration. GraphPad Prism 9 and using the [Agonist] vs. Response – Find ECanything nonlinear fit. Fit parameter F was constrained to 50, Bottom was constrained to 4 (the negative control for the assay) and the Top was constrained to be the same for all data sets (cell viability should be equal for all data sets as polymer concentration approaches 0). **b,** $LC_{50}$ values for each polymer. A post-hoc Tukey correction for multiple comparisons was used to determine statistical significance between $LC_{50}$ values. Adjusted p-values are reported. All data is shown as mean ± s.e.

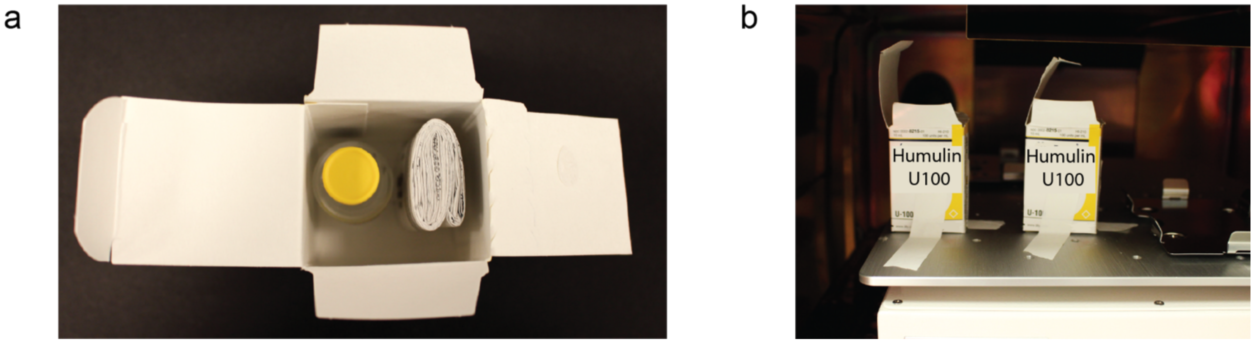


**Figure S7.** Photographs of relevant materials for high temperature aging study. **(A)** Cardboard packaging of 10 mL vial of Humulin including administration instructions. **(B)** Humulin boxes affixed to rotary shaker in an incubator.

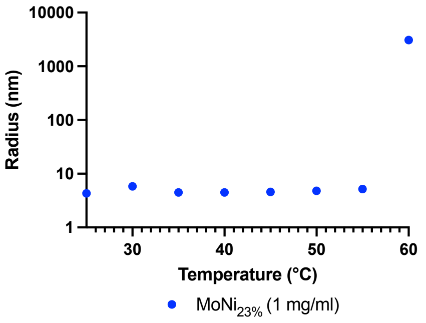


**Figure S8.** DLS (DynaPro Plate Reader II, Wyatt Technology) measurements of $MoNi_{23\%}$ at 1 mg/ml in miliQ water at temperature increments of 5 °C. A marked increase in radius is noted between 55 and 60 °C, an indication of the LCST of the polymer.

### SUPPLEMENTARY TABLES

#### Table S1 Molecular information for AC/DC Excipients

| Carrier | wt.% (Target) | wt.% by NMR (Experimental) | Dopant | wt.% (Target) | wt.% by NMR (Experimental) | $M_n$[a] (Da) | $M_w$[a] (Da) | Đ[a] |
|---|---|---|---|---|---|---|---|---|
| Morph | 77 | 74.5[b] | Nipam | 23 | 25.5[b] | 3200 | 3800 | 1.19 |
| Morph | 94 | 93.7[c] | Phe | 6 | 6.3[c] | 2900 | 3400 | 1.17 |
| Mpam | 92 | 91[d] | Phe | 8 | 9[d] | 5000 | 5400 | 1.08 |
| Morph | 100 | | - | | | 2300 | | 1.12 |

[a] Determined using Size Exclusion Chromatography calibrated using polyethylene glycol samples.
[b] Weight percentages difficult to determine due to overlapping spectra. Weight percentages estimated from post-precipitated NMR spectra by measuring the more resolved left half of the peak of Nipam (δ= 4.0, 0.5 H), doubling it, and subtracting it from the unresolved peaks of MORPH and Nipam (δ= 3.2-4.2, 7H (MORPH) 1H (Nipam)).
[c] Weight percentages calculated from post-precipitated NMR spectra of Morph (δ= 3.3-3.7, 8H) and Phe (δ= 7.6, 2H).
[d] Weight percentages calculated from post-precipitated NMR spectra of Mp (δ= 3.1-3.5, 7H) and Phe (δ= 7.6, 2H).
